# Amino acid properties, substitution rates, and the nearly neutral theory

**DOI:** 10.1101/2024.11.04.621833

**Authors:** Jennifer E. James, Martin Lascoux

## Abstract

Do the properties of amino acids affect their rates of substitution? The nearly neutral theory predicts that greater selective constraint leads to slower rates of evolution; similarly, we expect amino acids that are more different from each other to have lower rates of exchange because such changes are most likely to affect protein structure and function. Here we test these predictions, using substitution rates estimated from empirical amino acid exchangeability matrices. To measure degree of amino acid difference, we focussed on two physicochemical properties, hydrophobicity and molecular weight, uncorrelated metrics that are known to have important implications for protein structure and function. We find that for both hydrophobicity and molecular weight, amino acid pairs with large differences had lower rates of substitution. We also found that amino acids that differed in both properties had the lowest rates of substitution, suggesting that both physicochemical properties are under selective constraint. Mutation properties, such as the number of mutations or the number of transitions as opposed to transversions separating amino acid pairs, were also important predictors of substitution rates. The relationship between amino acid substitution rates and differences in their physiochemical properties holds across several taxonomically restricted datasets. This finding suggests that purifying selection affects amino acid substitution rates in a similar manner across taxonomic groups with different effective population sizes.

## Introduction

The neutral theory highlights the importance of purifying selection and conservation as forces in molecular evolution. Kimura stressed that more important proteins will evolve more slowly than less important proteins due to purifying selection removing variants that cause amino acid change (Kimura, 1983). This logic can easily be extended to variants that do cause amino acid change: under Kimura’s theory, we expect rates of exchange between amino acids that are dissimilar to be lower, as exchanges between more dissimilar amino acids are likely to cause large changes in protein structure and function. Investigating rates of exchange between different amino acid pairs therefore provides an interesting test of the neutral theory.

It is now clear that rates of substitution between different pairs of amino acids vary. For example, Smith (2003) defined amino acid substitutions as either ‘radical’ (physicochemically different) or ‘conservative’ (physicochemically similar) and found that radical substitution rates were lower than conservative rates for a dataset of *Drosophila* genes, which is evidence that radical substitutions are subject to stronger negative selection than conservative mutations. More recently, Weber and Whelan (2019) used several multi-species alignments to investigate radical and conservative substitution rates. While they found evidence that radical substitutions were removed by purifying selection more frequently than conservative substitutions in large populations, they did not find evidence for a difference in rates in species with small populations (mammals, birds and vertebrates). Similarly, Zou and Zhang (2019) estimated amino acid exchangeabilities from codon models between 90 pairs of closely related species, finding substantial variation across taxa. In contrast, Chen et al. (2019) used a dataset of nine pairs of species and found that substitution rates were strongly related across taxa rather than varying among species with different population sizes, and that rates were negatively correlated to the difference in physicochemical properties between amino acid pairs. There is therefore some contention as to whether amino acid substitution rates vary among taxonomic groups, or if they vary systematically with differences in amino acid physicochemical properties. There are also some methodological issues with past studies. For example, in studies that group amino acid substitutions as either ‘radical’ or ‘conservative’, it is not clear how radical or conservative substitution should be defined. Some amino acid substitutions are more ‘radical’ than others, as some involve exchanges between amino acids that differ in many properties.

There are also several methodological difficulties to estimating substitution rates. Firstly, a general requirement of many methods is a phylogenetic tree, which may differ across the genome due to incongruities between gene trees and the species tree, resulting in biases in substitution rate estimates (Carruthers et al., 2022; Mendes & Hahn, 2016). Secondly, accurate substitution rate estimation requires the careful alignment of orthologous genomic regions that must be either long enough or divergent enough to contain sufficient substitutions for inference without mutation saturation, i.e., the occurrence of multiple substitutions at the same site along the genome. Some judgement is therefore required as to whether a dataset is likely to provide estimates of substitution rates that are reasonably unbiased (Del Amparo et al., 2021). The problem of mutation saturation is expected to increase when evolution is rapid, or if the timescale under consideration is long. This has been demonstrated using datasets of viral genomes, which showed a decay of the transition:transversion ratio with time as expected if mutation saturation had occurred. There was a decay in the transition:transversion ratio with time even for datasets with short divergence times and small numbers of substitutions, in which there would be little *a priori* reason to expect mutation saturation (Duchêne et al., 2015).

An alternative approach is to consider exchangeability rates modelled at the amino acid level, rather than at the codon level. Amino acid substitution models are often estimated empirically from large datasets of multiple sequence alignments, with the resulting production of 20 by 20 matrices of exchangeability rates between amino acid pairs. Substitution matrices such as the Dayhoff, JTT, LG, and WAG matrices are widely used in phylogenetic analyses of protein sequences and are implemented in software tools including MEGA (Tamura et al., 2021), PAML (Yang, 2007) and RAxML (Stamatakis, 2014). Despite their wide use in phylogenetic methods, these models are often not considered in the context of the evolutionary insights they provide. These models can be computationally challenging to run and, due to the large number of parameters to be estimated, necessitate large datasets to prevent model over-fitting. However, recent advances in the field, including the development of user-friendly interfaces for estimating amino acid substitution models and the increasing availability of large sequence datasets, has made this approach increasingly tractable.

In this study, we used empirical models of amino acid substitution, incorporating recent advances in the field in order to address Kimura’s prediction that rates of exchange between more different amino acids will be lower. If rates of exchange between amino acid pairs are affected by the physicochemical differences between them, this suggests the action of purifying selection. We explicitly include mutational processes in our analyses, including GC content, the transition-transversion ratio, average amino acid frequencies, and number of mutational steps, to tease out the relative importance of mutational versus selective forces (Dagan et al., 2002). We also ensured that we accounted for amino acid composition when conducting our statistical analyses, an issue that has typically been overlooked in past studies. We investigated whether patterns in substitution rates differ across different timespans, or different taxonomic groups.

## Methods

We analysed amino acid exchange matrices calculated using QMaker (Minh et al., 2021) and nQMaker (Dang et al., 2022), implemented in IQ-TREE (Minh et al., 2020). Both QMaker and nQMaker estimate an amino acid exchange rate matrix using maximum likelihood, while also modelling variability in evolutionary rates across sites. The QMaker amino acid models of substitution rates that we used are estimated using a maximum likelihood procedure that makes three assumptions: that substitution rates are constant over time (homogeneous), that amino acid frequencies are at equilibrium (stationary), and that rates of exchange between two amino acids are equal, i.e., the rate of substitution from A to C is equal to that of C to A (reversible). nQMaker relaxes the time-reversibility assumption, producing time non-reversible matrices that do not obey detailed balance, that is, fluxes between amino acid pairs do not need to have the same magnitude.

We used the amino acid exchangeability matrices available online from IQ-TREE, as calculated over the Pfam dataset (version 31, El-gebali et al., 2019), and the matrices available for taxonomically restricted datasets (which included birds, insects, mammals, plants and yeast). To convert available exchangeability matrices to substitution rate matrices, we multiplied each exchangeability rate by the frequency of the appropriate amino acid in the dataset, converting the exchangeability rate matrix to the substitution rate matrix, *Q*. We use time non-reversible models for taxonomically restricted datasets, and time-reversible *Q* matrix in our analysis of the Pfam dataset, as previous studies showed that this model provides a better fit to these datasets (Dang et al., 2022; Minh et al., 2021). It is important to note that while the taxonomically restricted *Q* matrices were calculated using datasets of sequences present in their respective taxonomic group, most proteins are not specific to a particular clade, but instead are common to many species across the tree of life.

We took amino acid physicochemical properties from AAindex (Kawashima et al., 2008), an online database of amino acid indices, choosing properties of particular relevance to protein structure. We also considered a number of mutational amino acid properties. We calculated the average GC content over all codons per amino acid, and the average pyrimidine content over all codons per amino acid to approximate the transition-transversion ratio between codons. We also included the minimum number of mutational steps separating each amino acid. For all physicochemical and mutational amino acid properties, we calculated the absolute difference between amino acid pairs, creating a matrix of values per property. We then conducted statistical analyses on the data, after first log transforming amino acid substitution rates to better meet the assumptions of general linear models (Suppl. Fig. 1). We conducted general linear models including amino acid composition as random effects in our model, to account for the fact that amino acids appear multiple times among all pairs. We conducted all data manipulation and statistical analyses in R. All code and data tables used to conduct the analyses are available at: https://github.com/j-e-james/AminoAcidSubstitutions.

## Results

In this study, we investigated the relationship between amino acid substitution rates and differences in their physiochemical properties. It is not appropriate to consider all amino acid physicochemical properties together in statistical analyses, because many such properties are related; for example, polarity and hydrophobicity, and molecular weight and area (the correlation matrix between amino acid properties considered in this study is shown in Suppl. Fig. 1). We therefore considered a number of physicochemical properties, analysing the differences between amino acid pairs in their properties using a PCA (Fig. 1). Our PCA analysis split the amino acid physicochemical properties into charge/hydrophobicity associated metrics, and size-associated metrics: properties such as hydrophobicity, flexibility and contact energy, all of which are highly correlated (Suppl. Fig. 1), contributed to PC1 (40%), while size metrics such as maximum accessible solvent area, molecular weight and volume were the major contributors to PC2 (35%). There was little additional variance explained from further PCA axes, with PC1 and 2 together explaining 74% of the variance in the data. In our further analysis we therefore used these first two principal components as fixed effects in general linear models, with PC1 representing difference in charge and PC2 representing difference in size between amino acids.

**Fig. 1).**
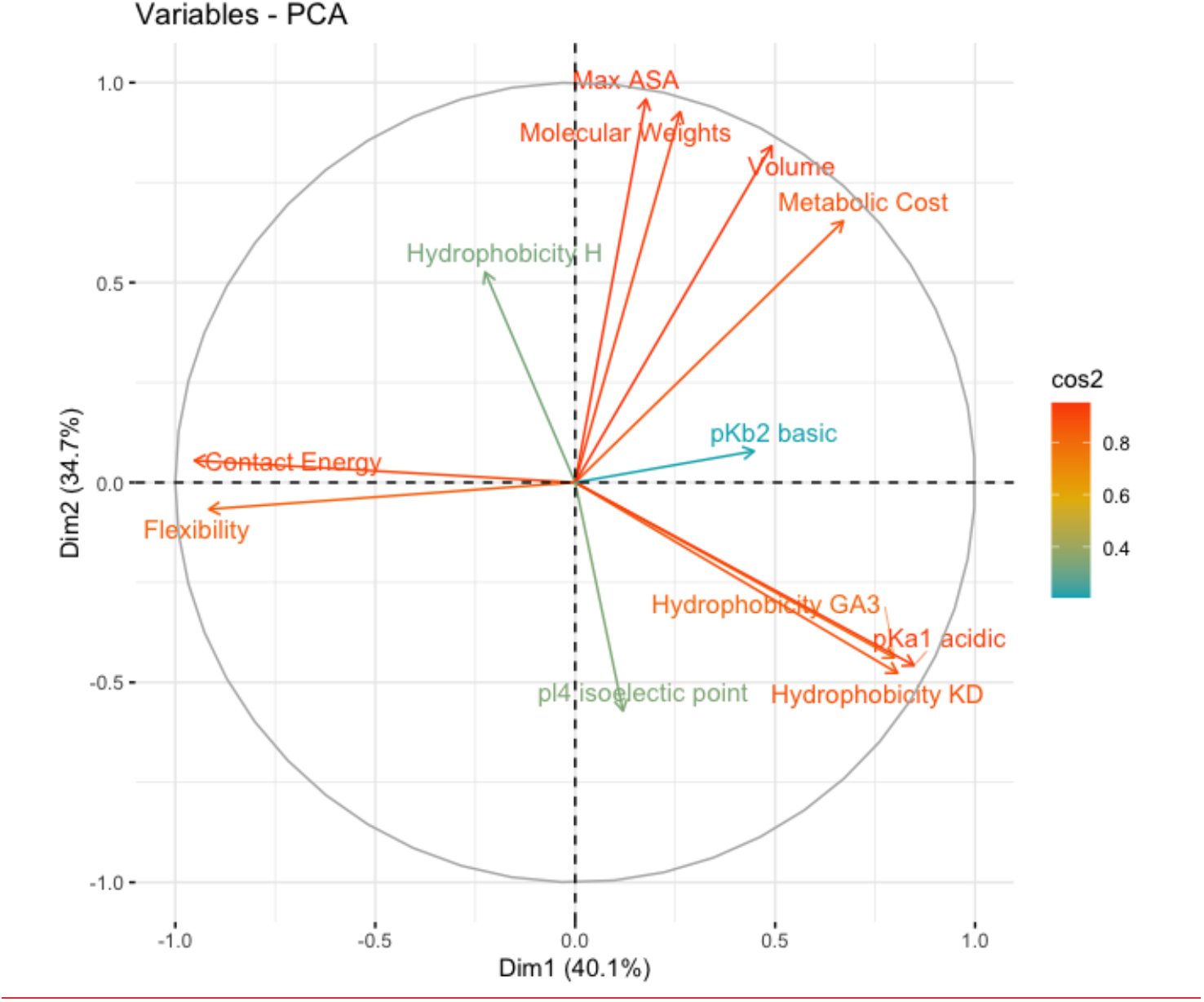
PCA of the difference in physicochemical properties between amino acid pairs, showing the variables and their contributions to the first two principle components. Variables are labelled on their respective arrows, with colours of text and arrows indicating cos^2^, the representation of the variable on the principle component.

We take as our estimates of exchangeability rates the non-time reversible *Q*-matrices from Minh et al. (2021), and the time reversible matrices from Dang et al. (2022). Such matrices reflect the structure of the genetic code, mutational biases, and selection pressure. We therefore conducted models taking into account the number of mutations that separate pairs of amino acids, the approximate transition to transversion ratio per amino acid, and the average difference in codon GC content between amino acid pairs.

We conducted linear mixed effect models of exchangeability rates, including amino acid composition as a random effect to account for the statistical non-independence in the data. We found that pairs of amino acids with large differences in physiochemical properties have low rates of exchangeability, and low substitution rates (Table 1). This fits the predictions of the nearly neutral theory: exchanges between amino acids with very different physiochemical properties are more likely to affect protein structure, and are therefore more likely to be removed by purifying selection, as opposed to exchanges between physicochemically similar amino acids. Both difference in amino acid size and difference in amino acid charge had a significant negative relationship with substitution rate (*p* < 2e-16 for both PC1 and PC2 in our model, Fig. 2), and also significantly interacted with each other, such that if the difference in one property was large, the relationship between the difference in the other property and exchangeability rate was more negative (an interaction term significantly improved model fit, *p* = 4e-14). This suggests that exchanges between amino acids that differ in both size and charge may be the most disruptive to protein function.

**Table 1.** Fixed effects of general linear model results. Shown is the best model, including an interaction term between PC1 and 2, and excluding difference in GC content.

|  | Estimate | Std.error | Degrees of freedom | t-value | *p* |
| --- | --- | --- | --- | --- | --- |
| Intercept | 0.076 | 0.084 | 105.00 | 0.90 | 0.373 |
| PC1, Charge | -0.436 | 0.021 | 342.94 | -20.46 | <2.0E-16*** |
| PC2, Size | -0.289 | 0.026 | 357.33 | -11.20 | <2.0E-16*** |
| Mutational steps slope | -0.422 | 0.026 | 350.50 | -16.15 | <2.0E-16*** |
| Transition:transversion difference slope | -0.202 | 0.086 | 371.25 | -2.36 | 0.019* |
| PC1: PC2 interaction | 0.083 | 0.011 | 341.77 | 7.86 | 4.9E-14*** |

**Fig. 2).**
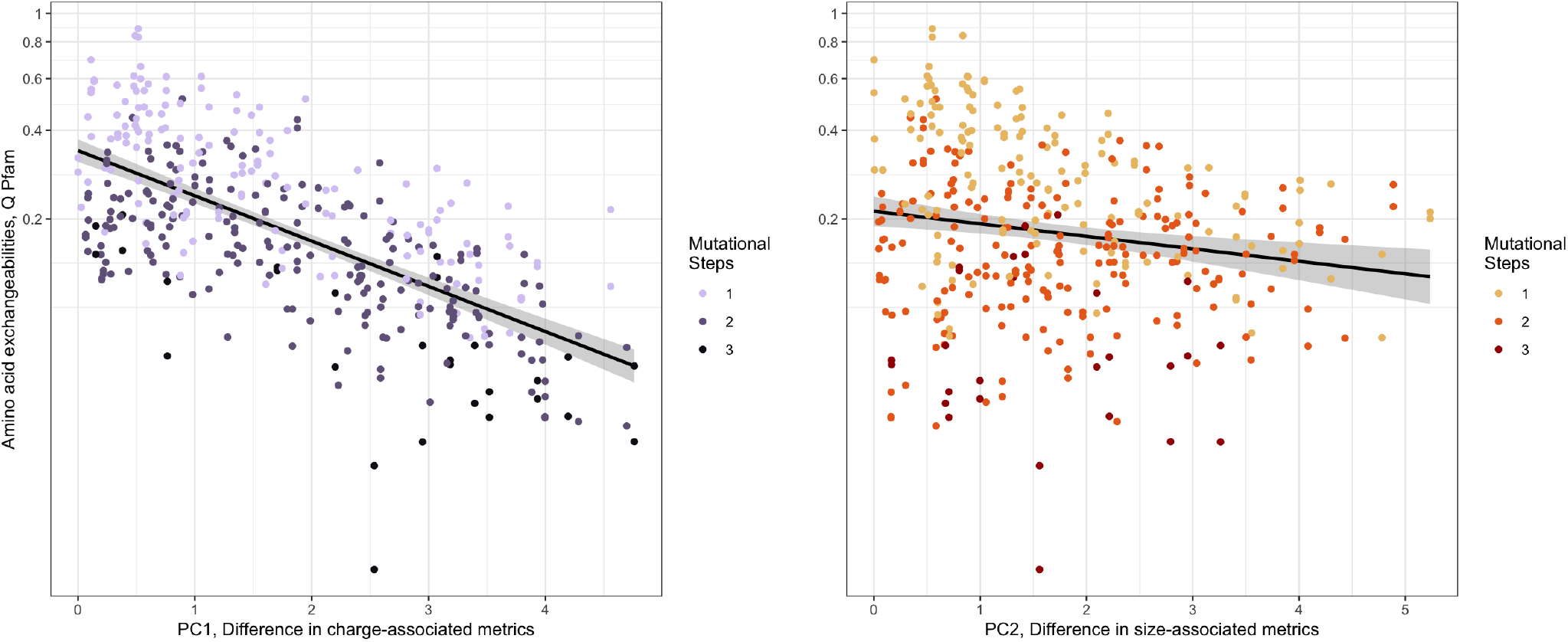
Relationship between amino acid differences in physicochemical properties (PC1, charge associated metrics, left, and PC2, size associated metrics, right), and substitution rates, for Pfam data. Points are coloured in a gradient from dark to light representing most to least mutations separating amino acid pairs. Black lines are to aid the eye only, as the plots show non-independent data. These lines are linear regression slopes for exchangeability rate ∼ difference in physiochemical property.

We found that average difference in codon GC composition does not affect the fit of our models (model comparison *p* = 0.40). Therefore, although local GC composition undoubtedly affects mutation rates and amino acid composition, it does not appear to influence amino acid exchangeability rates over long evolutionary timescales. In contrast, the number of mutations separating amino acid pairs significantly negatively impacts rates of exchange between them (shown in colour scales of Fig. 1, mutational steps in general linear model *p* < 2e-16). Similarly, amino acid pairs separated by more transversion mutations have lower exchangeability rates (average difference in purine and pyrimidine content in general linear model *p* = 0.019).

It is not surprising that amino acid exchanges that require multiple mutations occur at lower rates, and that this factor explains a considerable proportion of variance in our model (the proportion of variance explained by marginal effects including all parameters but GC is 0.68, while if mutational parameters are instead included as random effects, the proportion of variance explained by marginal effects drops to 0.35). We further investigated how the number of mutations separating amino acid pairs changes the relationship between rates of substitution and mutational and physicochemical differences by independently considering amino acids separated by different numbers of mutational steps. Even considering amino acid pairs separated by the same number of mutations, we find negative relationships between amino acid substitution rates and differences in their physiochemical properties (see Fig. 3 and Suppl. Table 1). For amino acids separated by one mutation, we find that physicochemical property differences are the primary driver of the relationship, with other mutational parameters not significantly improving model fit if included (AIC values with and without GC content and transition:transversion difference are 81.29 and 81.94 respectively, anova *p* = 0.098). This is particularly striking because amino acids that are more similar tend to be separated by only a single mutational step (Freeland & Hurst, 1998; Osawa & H., 1989), which we confirm is also true for the physicochemical properties considered in this study, although the difference is only significant when comparing our first principle component, associated with the difference in charge/hydrophobicity between pairs of amino acids (one/two mutations mean difference in PC1, charge: 1.55, 1.88, t-test *p* = 0.011, one/two mutations mean difference in PC2, size: 1.63, 1.72, t-test *p* = 0.5). For amino acid pairs separated by two mutations, the relationship between substitution rate and difference in charge and difference in size was negative and significant, with mutational parameters not improving model fit (*p* = 0.46), a very similar patten to that observed by amino acid separated by one mutation. For amino acid pairs separated by three mutations, only the negative relationship between substitution rate and difference in charge related metrics was statistically supported, likely due to the smaller size of this dataset. The primary effect that number of mutation steps has in our models is to change the intercept, such that the greater the number of mutation steps separating amino acid pairs, the lower the intercept (Fig. 2).

**Fig. 3).**
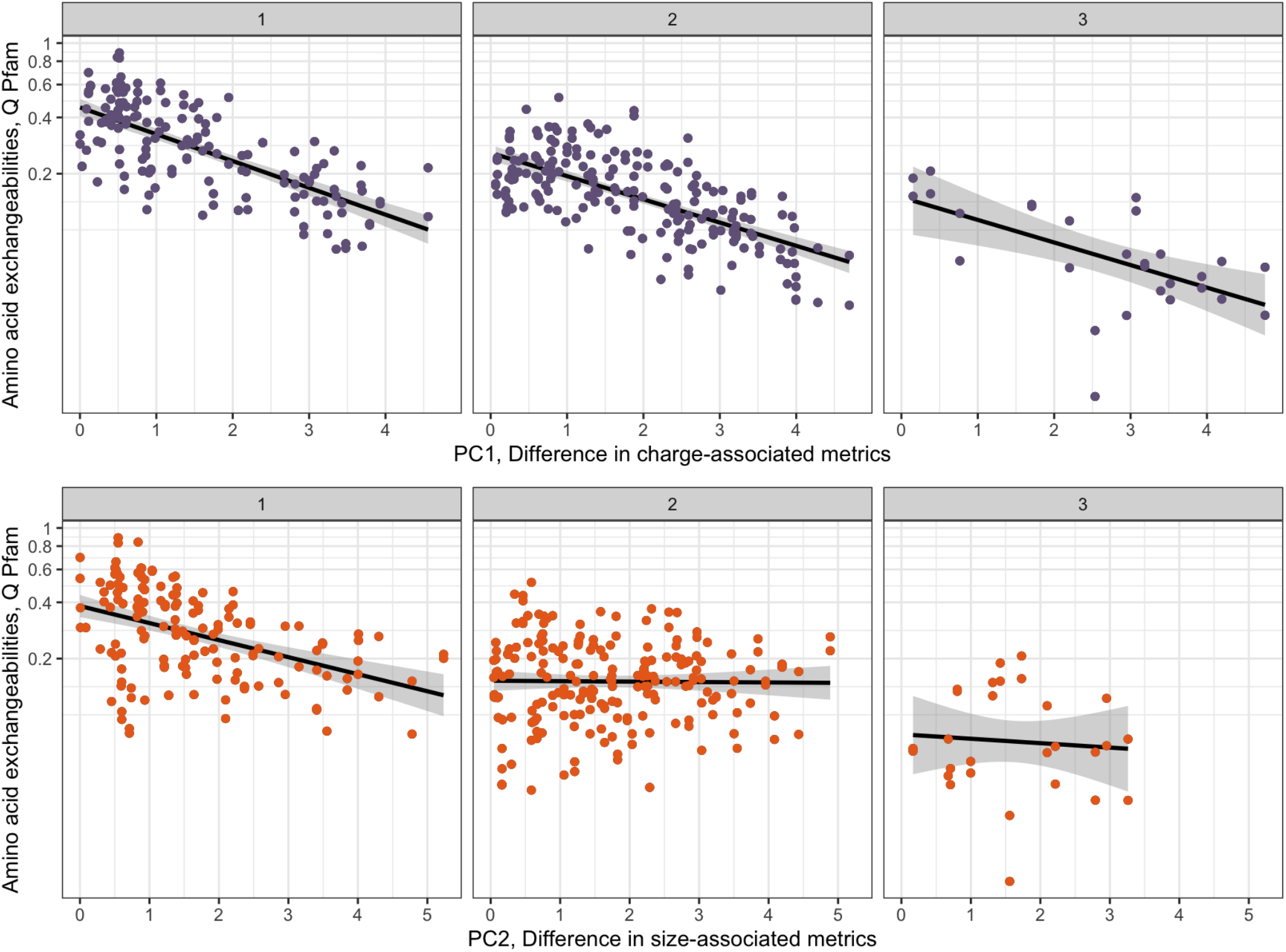
Relationship between amino acid differences in physicochemical properties (PC1, charge associated metrics, top, and PC2, size associated metrics, bottom), and substitution rates, for Pfam data. Data are separated into panels depending on the number of mutational steps (1, 2 or 3) between amino acid pairs. Black lines are to aid the eye only, as the plots show non-independent data. These lines are linear regression slopes for exchangeability rate ∼ difference in physiochemical property.

These results were calculated using the time-reversible exchange matrix for the Pfam dataset. They are therefore representative of long evolutionary timescales and are general across a whole range of species. We also repeated these analyses using more taxonomically restricted datasets, to investigate whether these patterns hold. We found results to be very similar when considering birds, insects, yeast, and mammal datasets, and when using exchangeability rates as estimated using time non-reversible matrices (model results shown in Suppl. Table 2). The relationship between exchangeability rates and the difference in these two physicochemical properties holds across all the datasets we tested. It is interesting that differences in amino acid physicochemical properties remain predictive of their substitution rates for both highly diverged sequences (e.g., the Pfam dataset) and for clade-specific datasets. Average difference in GC content was not related to exchangeability rate in any datasets, while number of mutational steps explained a large proportion of the variance in exchangeability rates across all datasets. We see some evidence that mutational factors may have a larger impact on amino acid substitution rates in small *N*_*e*_ groups than in large *N*_*e*_ groups, with mutational factors explaining a greater proportion of the variance in substitution rates in mammals, birds and plants than in yeast and insects (Suppl. Table 2). This may be because the rate of substitutions which require more than one mutational step are dependent on the rate of new mutations entering the population, which depends on census size.

## Discussion

Overall, we find a relationship between the difference in physicochemical properties between pairs of amino acids and the exchangeability rates between them, such that more different amino acids have lower rates of exchange. Additionally, while differences in amino acid exchangeability rates are known to exist across taxonomic groups, as evidenced by studies that find substantially better model fits for phylogenetic analyses when using taxon-specific exchangeability rate matrices (Del Amparo & Arenas, 2023; Minh et al., 2021), we find that a relationship between differences in physicochemical properties and substitution rates holds across all taxonomic groups considered. Our results suggest that, on average, there is evolutionary constraint acting at sites within proteins, such that changes with a small effect at the amino acid site level are more common. The exact shape of the relationship between amino acid substitution rates and their chemical differences has historically been under some dispute, with Kimura fitting an exponential curve to the data in his analysis (Kimura, 1983) while Gillespie, using more modern estimates of substitution rates and focussing on amino acids separated by a single mutation, found the polynomial distribution to be a better fit to the data, suggesting that the most frequent substitutions were between amino acids with small, but not the smallest, chemical differences (1991). Our findings are more similar to those of Kimura, and thus are in line with the nearly neutral explanation for this pattern: that non-conservative amino acid substitutions, which are more likely to disrupt protein structure and function, occur less frequently.

Our findings are a result of estimating amino acid substitution rates over many proteins and species. It is possible that there are biases or unconsidered factors in these datasets that could result in the observed relationship between amino acid physicochemical differences and substitution rates. For example, if particular pairs of amino acids are more often found in young proteins with high rates of turnover and high substitution rates, or if particular pairs of amino acids are often associated with rapidly evolving active sites, this would generate a relationship between the amino acid physicochemical differences and substitution rates, without the differences in physicochemical properties being causal. However, this is unlikely to be responsible for our results because we observe a trend with substitution rate for both charge/hydrophobicity associated metrics and size associated metrics, properties that are not correlated to each other (Suppl. Fig. 1), i.e., pairs of amino acids can be different in terms of charge/hydrophobicity but not size, and vice versa. Therefore, particular pairs of very exchangeable amino acids are probably not driving both trends.

Patterns of amino acid substitutions are determined by a combination of mutational forces, selection and drift, all of which may differ across species groups. In support of this, clade-specific differences in amino acid substitution rates have been found by a number of researchers. For example, Pandey and Braun (2020) estimated clade specific amino acid exchangeability rate matrices in order to investigate how much model variance could be explained by species clade. They found that, for the majority of proteins, the clade specific model fit was able to correctly assign proteins to their clade. Models of amino acid exchangeability have also been shown to cluster in terms of similarity in a manner akin to the branches of the tree of life, meaning that in general more closely related species have more similar amino acid exchangeability rate matrices (Scolaro & Braun, 2023). Likewise, we found that the two most closely related species sets in our datasets, mammals and birds, did have more similar amino acid exchangeability matrices, and more similar substitution rate matrices, than other species sets (Suppl.Tables 1 and 2). Further development of time non-reversible and taxon-specific models of amino acid substitution may shed light on the biological underpinnings of the observed variation in substitution patterns across taxa.

Given our specific nearly neutral hypothesis for the observed relationship between amino acid substitution rates and differences in their physicochemical properties, we might also expect specific differences among clades in terms of this relationship. In species with large effective population sizes, we might expect substitution rates among amino acid pairs that differ in their physicochemical properties to be even lower than in species with small effective population sizes due to stronger purifying selection, resulting in a steeper relationship between substitution rates and physicochemical property differences.

This pattern was observed in some previous studies (Weber & Whelan, 2019; Zou & Zhang, 2019); however, it is not particularly apparent in our results. The relationship between substitution rates and the physicochemical properties of amino acids was similar among datasets, despite some differences in substitution rate matrices among taxonomic groups. It could be that, even for those groups in our dataset with small effective population sizes, selection against large changes in the physicochemical properties of amino acids is sufficiently strong on proteins overall that genetic drift does not have a large effect on substitution rates. It may be that *N*_*e*_ is sufficiently large across all studied taxa that the effect of further increasing *N*_*e*_ is small.

The further refining of amino acid substitution models is an important area of future research. In particular, while variation in substitution rates across sites is known to exist and is modelled when estimating amino acid exchangeabilities, it is possible that not only rates but also patterns of substitution vary across sites within proteins. For example, while solvent exposed sites are known to evolve more rapidly (Goldman et al., 1998), Pandey and Braun (2020) found that patterns of substitution also vary on a finer scale across such sites, after estimating amino acid substitution models for exposed and buried residues separately. The influence of protein structure on patterns of molecular evolution was also explored by Perron et al. (2019), who, using protein sequences for which structural data was available, estimated substitution models while also incorporating information on the rotational configuration of amino acid side chains. The resulting models had an expanded number of possible exchangeabilities, as each amino acid could be exchanged not only with every other amino acid, but also one of three side chain conformational states, expanding the 20 by 20 amino acid exchangeability matrix to a 55 by 55 exchangeability matrix. The models performed well when fitted to both real and simulated data and resulted in biological insight: the authors found evidence that side chain configuration was often conserved when an amino acid exchange occurred, and that highly exchangeable amino acids often had similar side chain geometries. More recently, Ferreiro et al. (2024) implemented an ABC approach to model substitutions while incorporating information on the likely selective constraints acting on proteins (including amino acid physicochemical properties), and allowing for site-dependent evolution. Our results, and the ongoing developments of such methods, highlight the importance of incorporating knowledge of amino acid properties and protein structure into models of molecular evolution.

## Acknowledgements

This work was supported by a SCAS Natural Sciences Fellowship and by a grant from the Wenner-Gren Foundation. This work was also supported by the SciLifeLab & Wallenberg Data Driven Life Science Program (grant: KAW 2024.0159).

